# Supervised and Unsupervised Classification of lncRNA Subtypes

**DOI:** 10.1101/2020.07.20.211433

**Authors:** Rituparno Sen, Jörg Fallmann, Maria Emília M. T. Walter, Peter F. Stadler

**Affiliations:** Bioinformatics Group, Department of Computer Science, and Interdisciplinary Center for Bioinformatics, University of Leipzig, Härtelstraße 16-18, D-04107 Leipzig, Germany; Departamento de Ciência da Computação, Instituto de Ciências Exatas, Universidade de Brasília; German Centre for Integrative Biodiversity Research (iDiv) Halle-Jena-Leipzig, Competence Center for Scalable Data Services and Solutions, and Leipzig Research Center for Civilization Diseases, University Leipzig, Germany; Max Planck Institute for Mathematics in the Sciences, Inselstraße 22, D-04103 Leipzig, Germany; Institute for Theoretical Chemistry, University of Vienna, Währingerstraße 17, A-1090 Wien, Austria; Facultad de Ciencias, Universidad National de Colombia, Sede Bogotá, Colombia; Santa Fe Institute, 1399 Hyde Park Rd., Santa Fe, NM 87501

**Keywords:** host gene, miRNA, snoRNA, *k*-mers, secondary structure, random forest

## Abstract

Many small nucleolar RNAs and many of the hairpin precursors of miRNAs are processed from long non-protein-coding (lncRNA) host genes. In contrast to their highly conserved and heavily structured payload, the host genes feature poorly conserved sequences. Nevertheless there is mounting evidence that the host genes have biological functions. No obvious connections between the function of the host genes and the function of their payloads have been reported. Here we inverstigate whether there is an association of host gene function or mechanisms with the type of payload. To assess this hypothesis we test whether the miRNA host genes (MIRHGs), snoRNA host genes (SNHGs), and other lncRNAs host genes can be distinguished based on sequence and structure features. A positive answer would imply a correlation between host genes and their payload. While the three classes can be distinguished reliably when the classifier is allowed to extract features from the payloads, this is no longer the case when only sequence and structure of parts of the host gene distal from the snoRNAs or miRNA payload is used for classification. Our data indicate that the functions of MIRHGs and SNHGs are largely independent of the functions of their payloads. Furthermore, there is no evidence that the MIRHGs and SNHGs form coherent classes of long non-coding RNAs distinguished by features other than their payloads.

## 1. Introduction

A wide variety of molecular and biological functions have been reported for long non-coding RNAs (lncRNAs), recently reviewed in [1]. Specific lncRNAs regulate chromosome architecture and chromatin remodeling, modulate inter- and intrachromosomal interactions and recruit or prevent the recruitment of chromatin modifiers. LncRNAs regulate transcription by forming R-loop thus recruiting transcription factors and interfere with the Pol II machinery to inhibit transcription. Other lncRNAs are structural components on nuclear bodies. In the cytoplasm, lncRNAs regulate mRNA turnover, translation and post-translational modification. Many lncRNAs serve as host genes for the production of small RNAs [2], including microRNAs (miRNAs) and small nucleolar RNAs (snoRNAs), or act as sponges that modify the miRNA pool [3]. In contrast to protein-coding genes, where function is closely tied to protein families, sequence similarity is a poor predictor of functional similarity in lncRNAs. As a consequence, it has remained impossible to predict the biological function or molecular mechanism of lncRNAs for sequence data alone. This is in stark contrast also to the small, structured ncRNAs. These are readily recognized through their highly conserved sequences (such as ribosomal or spliceosomal RNAs), or by class specific features (such as the cloverleaf shape of tRNAs or the ultra-stable hairpins of miRNAs) [4].

Unsupervised clustering using normalized *k*-mer abundances as similarity measure revealed an association of *k*-mer profiles with lncRNA function, in particular with protein binding and sub-cellular localization [5]. With a plethora of RNA-binding proteins typically recognizing a wide array of local binding motives that may be structured, modular, and gapped [6], the correlation of short *k*-mers and function is not surprising. Nevertheless, it remains unclear whether there are distinct, well-separated classes of lncRNAs or whether the universe of lncRNAs is organized as a continuum of functions and associated molecular features.

Here we consider two classes of lncRNAs that, in addition to any intrinsic function they might have, also act as host genes for small RNAs with well known and quite specific molecular functions: snoRNA hosts genes (SNHGs) and miRNA host genes (MIRHGs). Due to the differences in their payloads (i.e., the snoRNA/miRNA part of the transcript), they undergo distinctive processing.

The processing of snoRNAs for SNHGs in human is linked to splicing, with snoRNAs exclusively located in introns, see e.g. [7]. SNHGs have received rapidly increasing attention in particular in cancer research, see e.g. [8–13] and the references therein. SNHG15, for instance, is dysregulated in a wide variety of cancers and interacts with several distinct molecular pathways apparently using different molecular mechanisms, reviewed in [10]. Similar, there are several competing, not mutually exclusive, mechanistic explanations for the function of GAS5 [8]. There is strong indication that at least one of the modes of action of most SNHGs is to act as miRNA sponge [11,14]. The snoRNAs themselves are mostly involved in the maturation of rRNAs and snRNAs, where they direct chemical modifications, although an increasing number of secondary functions have been reported very recently [15]. However these do not seem to be coupled to functions of the host genes.

The precursors of canonical miRNAs, whether intronic or exonic, are extracted from the primary transcript by the microprocessor complex centered around Drosha and DGCR8 [16]. A second smaller group of miRNAs, the mirtrons, are processed with the help of the splicing machinery [17]. Functionally, miRNAs orchestrate post-transcriptional gene silencing, affecting a large fraction of protein-coding transcriptome [18,19]. In contrast to SNHGs, very few MIRHGs have known functions independent of the miRNAs that they harbor. Two notable examples are MIR100HG and MIR31HG [20]. MIR100HG has been reported to interact with HuR/ELAVL1 [21] and to form RNA-DNA triplex structures with the p27 locus [22], and possibly it may also be a ceRNA [23].

In this contribution we ask a simple question related to the global map of the lncRNA universe proposed in [5]: are SNHGs and MIRHGs distinct classes of lncRNA? To answer this question we use supervised and unsupervised methods of machine learning and ask whether machines can be trained that reliably separate SNHGs and MIRHGs from each other and from a background set of lncRNAs that harbor neither snoRNAs nor miRNAs, which we will refer to as NoHGs.

In order to identify which parts of the sequences are informative on the RNA class, we devise four different collections of subsequences (see Fig. 1) and assay to what extent they can distinguish between the tree RNA classes by a supervised machine learning approach. In the first test, the payloads are covered. Since miRNA and snoRNAs have been shown to be easily distinguished by machine learning methods (see e.g. [24–27] and the references therein), it is no surprise that the three RNA classes are distinguishable from this information. We then consider the sequences flanking the payload to test whether it carries distinctive information on the mode of processing; as we shall see, the results are discouraging. Somewhat surprisingly, however, the classification accuracy increases significantly, when instead of the often intronic flanking sequence the flanking exonic sequence is used. Using sequences far away from the payload to assess payload-indenpendent properties, finally, we observe very poor classification. Unsupervised methods, furthermore, also fail to separate the three lncRNA classes. In the following section we describe these observations in detail and discuss the impact of particular types of features. The implications are discussed in Section 3.

**Figure 1.**
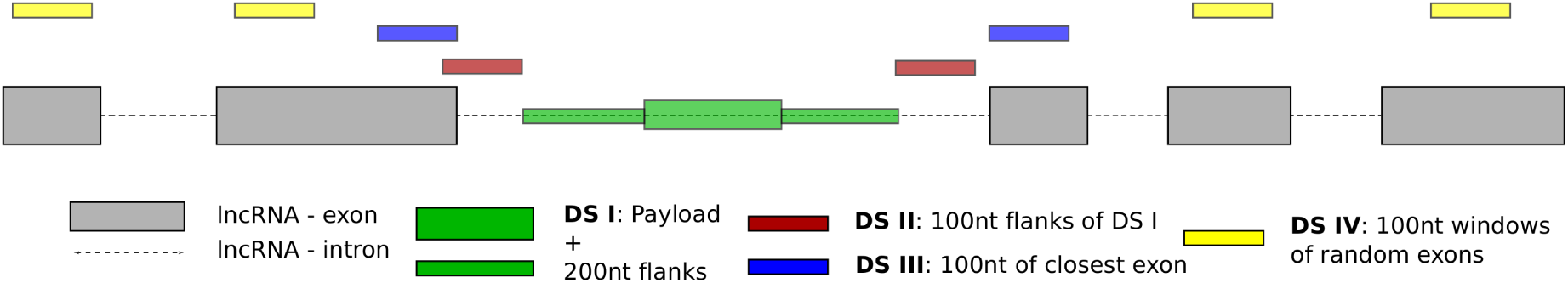
A schematic of the datasets curated for this study and their distribution over the gene body of a generic host-lncRNA. DS I (green) consists of the payload and 200nt flanking sequence. DS II (red) flanks DS-I by 100nts. DS III consists of the first 100nts of the exon closest to the annotated payload. DS IV consists of non-overlapping 100nt windows taken from random exons of the host-lncRNA. More details can be found in section 4.1

## 2. Results

To answer the underlying question of whether or not SNHGs and MIRHGs are indeed distinct classes of lncRNAs, we ask to what extent they can be distinguished from each each other and from a NoHG by mean of machine learning techniques. Since it is well-known that the payloads (miRNAs and snoRNAs) can be distinguished from each other and from background RNA in this manner [24–27], the interesting research question is whether this remains true when payload-related information is withheld from the classification machinery. To this end we carefully curated four distinct data sets covering different levels of potentially payload associated information. The structure of the curated datasets is summarized in figure 1 and table 2 in the Methods section. These are evaluated in both supervised and unsupervised settings.

**Table 1.**
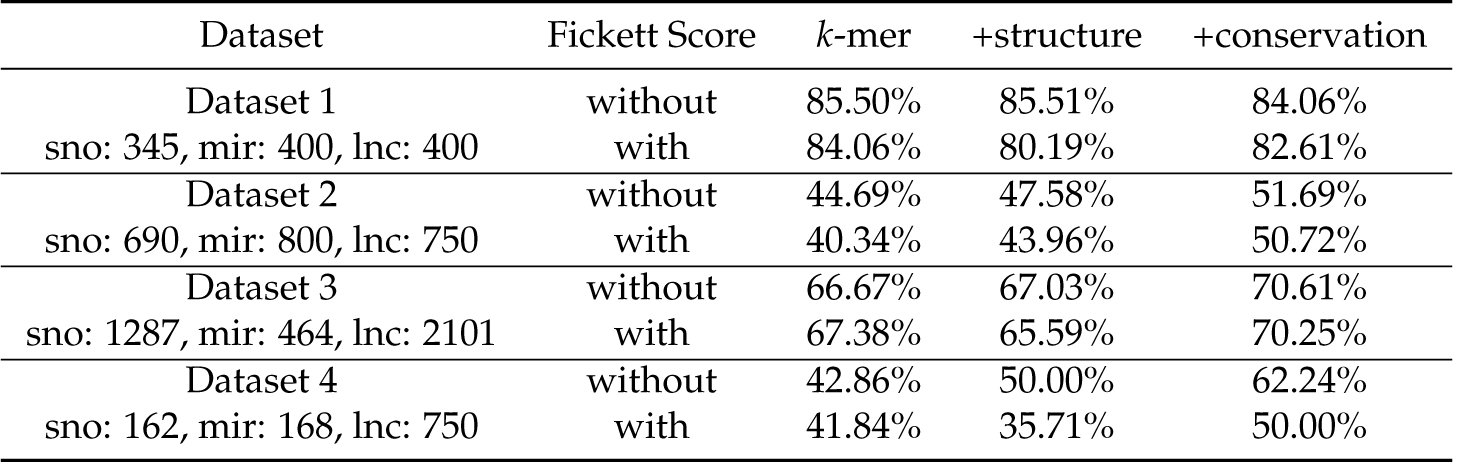
Overview of 10-fold cross validation accuracy for supervised machine learning on combinations of data and feature sets (*k*-mers only, *k*-mers plus secondary structure, or *k*-mers plus secondary structure and sequence conservation), both with and without the Fickett score as measure of coding potential.

| Dataset | Fickett Score | $k$-mer | +structure | +conservation |
| --- | --- | --- | --- | --- |
| Dataset 1 | without | 85.50% | 85.51% | 84.06% |
| sno: 345, mir: 400, lnc: 400 | with | 84.06% | 80.19% | 82.61% |
| Dataset 2 | without | 44.69% | 47.58% | 51.69% |
| sno: 690, mir: 800, lnc: 750 | with | 40.34% | 43.96% | 50.72% |
| Dataset 3 | without | 66.67% | 67.03% | 70.61% |
| sno: 1287, mir: 464, lnc: 2101 | with | 67.38% | 65.59% | 70.25% |
| Dataset 4 | without | 42.86% | 50.00% | 62.24% |
| sno: 162, mir: 168, lnc: 750 | with | 41.84% | 35.71% | 50.00% |

**Table 2.** Distribution of sequences contained in each dataset.

|  | Dataset 1 | Dataset 2 | Dataset 3 | Dataset 4 |
| --- | --- | --- | --- | --- |
| sno | 345 | 690 | 1287 | 162 |
| mir | 400 | 800 | 464 | 168 |
| lnc | 400 | 750 | 2101 | 750 |

### 2.1. Supervised Machine Learning

In order to gain insight into the usefulness of selected features, we trained a set of random forest classifiers with combinations of distinct feature sets extracted from different genic regions of the three lncRNA classes. Details on this procedure can be found in 4.1, results of 10-fold cross-validation (10xCV) on all data and feature sets can be found in table 1, details to training, testing and CV can be found in section 4.4.

#### Payload-aware prediction

Input data that include the full payload sequences are expected to allow the recognition of the payload type and the absence of a payload, respectively. The classification task on such data therefore serves as benchmark for our dataset/feature-selection strategy and expect that the three classes of lncRNAs can be distinguished with satisfactory reliability on *Dataset 1*, which makes the payload, i.e., miRNA and snoRNA precursors, resp., explicitly available to the classifier. We trained a random forest classifier, with feature sets ranging from sequence only features like weighted *k*-mer sets to structuredness and conservation (see section 4.3). Ficket score was added as extra feature to each comparison, to investigate the influence of coding potential. Table 1 summarizes the results for 10xCV on a training/test set split of 80/20. Accuracy of > 84% shows that random forests are indeed suitable for this kind of classification task. An interesting finding, however, is that the inclusion of information derived from structure/conservation features leads to a decrease in accuracy, probably due to the high structuredness of both miRNA and snoRNA payloads and similar conservation patterns. We note, finally, that a perfect separation of snoRNAs and miRNA precursors cannot be expected since there is a group of ncRNAs that appear to be in transition between the two functions [28–30].

#### Influence of the payload

The payload potentially, and in the case of MIRHGs even likely contains sequence or structure level signals needed for processing. We thus assume that machine learning techniques will be able to pick up on those signals. To test whether the actual payload of the host genes is necessary, or whether processing-related features in the adjacent sequence are sufficient to classify the host-gene type, we restricted *Dataset 2* to 200 nucleotides of flanking sequence on either side of the payload without overlap and random lncRNA subsequence for NoHGs. We observe that the exclusion of actual payload sequences had a strong impact on prediction accuracy, dropping from > 84% to below 45%. Although integration of structure and conservation features led to an increase of accuracy to more than 51%, it becomes clear that the sequences in close vicinity to the actual payload are insufficient to reliably identify the type of payload. Ranking of features according to their importance for classification shows that with the exclusion of the actual payload, sequence and conservation become important features for the classification task (supplementary table 2).

#### Exonic sequences only

Since snoRNAs are located in introns of the host gene, *Dataset 2* is almost exclusively restricted to intronic regions. However, functions of the matured host genes or other lncRNAs presumably reside in their exonic sequences. Furthermore, sequence and structure features involved in splicing as well as the initial processing of exonic miRNAs are likely to be found in the matured transcript. *Dataset 3*, which is designed to focus on such effects, comprises the 200nt of exonic sequence flanking the payload-bearing intron in the case of an intronic payload or the miRNA precursor in case of exonic miRNAs. Although by far not as good as the accuracy for classification including payloads, we see accuracy rises again to above 66%, including structure and conservation features even > 70%. We conclude that the exonic flanking regions contain more reliable information related to the processing of payload than the flanking sequences in the primary sequence. For miRNAs this can be explained in particular by sequence motifs associated with microprocessor activity [31,32]. For snoRNAs, their obligatory location in (usually short) introns also supports a connection with splicing, see also [7,33].

#### Random exonic sequences

Based on results from analysis of *Datasets 1-3*, we infer that exonic regions flanking the payload contain some amount of information about the type of the payload. Hence, we decided to test this theory, curating *Dataset 4* from random portions of exonic regions around the payload. Although exonic, these regions should be less prone to contain information specific to the payload compared to those in *Dataset 3*. Although the accuracy after inclusion of structure and conservation is higher than for payload flanking sequences of *Dataset 2* (> 62% in comparison to > 51%), it is still well below the accuracy obtainded from the flanking exonic regions with > 70%, supporting our hypothesis.

#### Fickett score

Invented as a measure for coding potential and a go-to feature for separating coding from non-coding RNAs by means of ML [34], we investigated whether the Fickett score [35] can be used as informative feature for our classification problem as well. While we observed a small increase in accuracy for flanking exons (*Dataset 3*) when only sequence features are used, every other prediction did not profit from its inclusion, thus the Fickett score does not constitute a meaningful feature for the classification problem at hand.

#### RNA Secondary Structure

SnoRNAs are highly structured RNA species, depending on the correct fold for their biological function. MiRNAs on the other hand are usually loaded into the RISC complex as single strands, for prior processing however, they are also heavily structured. Thus, secondary structuredness, or the lack thereof, could indicate regions which are better suited for the integration of either payload. Using RNAplFold [36] of the ViennaRNA package [37,38], we included the probability of being unpaired as feature vectors into our predictions, as explained in detail in section 4.1. In contrast to the Fickett score, inclusion of these features always increased accuracy, although not by much in most cases. In combination with Fickett Score, prediction accuracy is even decreased, except for dataset 2. Taken together, secondary structure features are informative for the classification task at hand, although their impact appears to be smaller than one might have expected, probably because (conserved) secondary structures are one the one hand abundant across the transcriptome and on the other hand do not seem to be characteristic for lncRNAs [39].

#### Sequence Conservation

Sequence conservation is a key indicator of biological function. It therefore seems sensible to include conservation features in our classifications. However, this limits application to genomic regions where reliable sequence alignments can be constructed, from which conservation scores can be deducted. We integrated the PhastCons score, i.e., the probability that a given position is part of a conserved region [40], as a convenient measure of position-wise conservation, see section 4.1. Similar to structure derived features, adding conservation increases accuracy except when focusing directly on the payload. This effect is strongest when random exonic regions (*Dataset 4*) are used and could hint towards a general trend of conservation differences between the three classes of lncRNAs investigated.

### 2.2. Unsupervised machine learning

So far, we have focused on identification of useful features for classification, comparing different datasets and feature combinations via means of random forest based supervised ML. Results show a significant decrease in accuracy for all approaches excluding the actual payload. Recent findings by Kirk *et al*. [5] *et al*. present a link between *k*-mer profiles and lncRNA function that could potentially also link to the payload of lncRNA host genes. Thus, we next focused on unsupervised clustering approaches. An initial principal component analysis (PCA) of the four data sets did not reveal a credible clustering or separation between NoHG, MIRHG, and SNHG, see supplemental figures S5-S8. Furthermore, using *k*-means clustering we found the assignment of the three groups of lncRNAs to the clusters is effectively random: accuracies for all combinations of features and datasets are below 36% (see supplemental table S1 and supplemental figures S1-S4). Also the use of a convolutional neural network (CNN) with an autoencoder showed no credible separation of the lncRNA classes, which is to be expected, however, given the small size of training and test sets. Unsupervised clustering methods appear to be unable to classify lncRNAs by their RNA payload, at least with the data available at present. It appears that the signal that can be obtained from the small miRNAs or snoRNAs is drowned out by the differences among the much larger hosts. This is at least indicative that there are no strong features that identify MIRHGs or SNHGs as coherent subgroups in lncRNAs.

### 2.3. MicroRNA target sites as a feature

At least some SNHGs can act as miRNA sponges [11,14]. If this is the primary function of the exonic part of SNHGs, these lncRNAs should be recognizable using the distribution of miRNA binding sites as features for classification. To test this hypothesis, we retrieved predicted and experimentally validated miRNA binding sites from miRTarBase [41], which in contrast to other sources contain not only 3’UTR located sites but any experimentally validated binding site regardless of its genic location. Intersection of this resource with our list of SNHGs and MIRHGs, however, revealed no significant overlap of miRNA binding sites and any of our host lncRNAs. To improve on this list of experimentally validated targets in regards to lncRNAs, which constitute less than 0.1% of the reported target genes, we used the *k*-mer feature to scan for miRNA seed region matches. A list of seed regions for each miRNA is readily available at TargetScan [42]. We restricted our search to regions covered in the analyzed datasets for which we had weighted *k*-mers available. No conclusive enrichment for any of the analyzed seeds could be detected comparing SNHGs and the other classes. This approach is of course limited due to the rather small regions we cover in our datasets, for a comprehensive analysis the whole gene body of all host genes has to be taken into consideration. Such an approach, however, is not feasible in context of our machine learning based classification task.

## 3. Discussion

In this contribution we asked whether there is a clear distinction between the host genes of snoRNAs and miRNAs on the one hand and lncRNAs without such highly conserved payloads on the other hand. Our answer is largely negative. While ML methods readily distinguish the three classes based on generic features of their payload, the classification task appears to become very difficult if the information about the payload and its immediate vicinity is not made available to the classifier. While the immediate vicinity – both exonic and intronic – appears to contain payload-specific information, the association between payload and features obtained from distant regions of the lncRNA is weak. Features derived from structuredness and conservation of these regions have in all cases a positive effect on classification accuracy and are readily ranked among the most important features. They do, however, not compensate for the weak association.

Given that there is mounting evidence for biological function of not only lncRNAs but also specifically for snoRNA and miRNA host genes, we argue that this *lack of detectable association* is of biological interest. It suggests that the function of the host genes is not closely tied to the function of the payload. This is in stark contrast to the protein-coding host genes of snoRNAs, many of which encode ribosomal proteins [43] and thus also contribute to the maturation of the ribosome.

Although a function as miRNA sponge has been reported for many SNHGs, we could not detect features that might connect the sequence or structure of the SNHGs with specific *k*-mers (namely those complementary to the seed regions of miRNAs) or to predicted miRNA target sites. Considering available, experimentally validated miRNA targets limits such a study for ncRNAs in general and even more so for lncRNAs, as they represent only a very small fraction of known target genes. A dedicated investigation into that matter is missing and would help to shed light onto the regulatory interplay between SNHGs, MIRHGs and their payload.

Applying *k*-means clustering to our datasets and features showed poor classification accuracy. Even though lncRNAs as such may harbor sequence motifs that give away conserved RNA binding protein target function among lncRNA groups, our investigation shows only poor classification potential for MIRHGs and SNHGs, even if the payload is considered. Although no distinctive clusters of MIRHGs and SNHGs were observed, our data are consistent with clustering of lncRNA classes observed in [5]. There, prominent SNHGs such as GAS5 have remained outside the identifiable clusters.

From an evolutionary point of view it may not be surprising that the host genes of miRNAs and snoRNAs do not exhibit recognizable class-specific features. Most likely, the molecular function of the payload, miRNAs or snoRNAs is much older and pre-dates functions of the non-coding host genes, which likely arose secondarily, maybe long after the transcripts have come under negative selection as host genes. The lack of common features of the host genes together with the usually very poor sequence conservation suggests they may even have acquired different functions in different lineages. A better understanding of the host genes thus will require a much more detailed investigation into the patterns of conservation than what is available at present. With rise of new sequencing technologies and advances in functional screening methods we can expect that more detailed data on host gene functions will be forthcoming. This may revise the current picture of distribution of lncRNA functions, which shows only a rather loose association of biological function and molecular mechanism with sequence and structure features of transcripts.

## 4. Materials and Methods

### 4.1. Sequence retrieval

For a comprehensive investigation into the separability of SNHGs, MIRHGs, and NoHGs we first had to curate data from available annotation and define sets of features than can be reliably used for training and prediction. MiRNA sequences were collected from miRBase [44], snoRNA sequences from the Human snoRNA Atlas [45] and snoDB [46], and lncRNA sequences were retrieved from GENCODE v29 and GENCODE v33 [47]. Where multiple lncRNA isoforms were annotated we used the longest one as representative. The coordinates of miRNA precursors from miRBase were used to identify MIRHGs and the exact position of the payload(s) within the MIRHG. All overlaps were computed with the bedtools suite [48] and customized Python scripts. Since all snoRNAs are intronic, no further processing was needed for SNHGs. In MIRHGs we defined both intronic and exonic miRNA precursors with 100 nt flanking sequence as the “payload”. The intron/exon annotation provided by the GENCODE annotation was used to define the exonic part of the lncRNA.

### 4.2. Datasets

The four datasets differ in regions selected, e.g. including payload, i.e., miRNAs and snoRNAs, or not. The difference is illustrated in Fig. 1. All data used for classification were extracted following GENCODE annotation (version 33) using custom scripts and the bedtools suite [48]. For purpose of training and testing, datasets were always balanced based on the smallest number of sequences available for any given class, to prevent prediction artefacts.

#### Dataset 1

For a set of 400 MIRHGs and 345 SNHGs, we extracted the payload sequence together with 100nt flanks. Flanks are considered to contain information necessary for the processing of the payload, *i. e*. for example processing of pre-miRNAs by the DROSHA/DICER machinery. As negative control set we collect 400 subsequences of random lncRNAs excluding overlaps with our positive sets on gene and sequence and level. Sequence length was set to 500nt as this was the mean length of sequences in our positive sets.

#### Dataset 2

To investigate the influence of the information derived directly from payload on our classification approach, we next prepared a dataset consisting of 800 samples from MIRHGs, 690 samples from SNHGs and 750 samples from NoHGs. For the host genes we extracted 100nt windows flanking the regions used as dataset 1. Overlaps between extracted sequences were discarded. These can arise from closely spaces payloads. This makes sure we have no information on the actual payload while we preserve close-by context. Here we do not focus on the genic region of these flanks. As negative set we again derive random subsequences of lncRNAs without overlap with the positive set, this time of size 100nt to match the positive sequences.

#### Dataset 3

Since the genic sequence information itself is of particular interest, we constructed a set that is similar to dataset 2, with the additional constraint that 100nt sequences are chosen from the closest exon up- and downstream of the payload. Dataset comprises 464 miRNA and 1287 snoRNA host gene sequences and differs from dataset 1 by exclude payload as well as the intronic sequence surrounding the payload. As negative set we selected at random 2101 exonic regions of length 100 from randomly picked lncRNAs without overlap to the positive set.

#### Dataset 4

To avoid any localized effect associated with the payload, we constructed dataset 4 by selecting multiple, non-overlapping regions of length 100nt from the exonic parts of the host genes. This data set contains 168 sequences extracted from miRNA precursors and 162 extracted sequences from snoRNA precursors. For the negative set we chose multiple, non-overlapping, 100nt windows of random exons of random lncRNA genes without overlap with the positive set, amounting to 750 sequences.

### 4.3. Features

The following features were computed for each sequences in the four datasets described above:

1. *k*-mer counts for 2 ≤ *k* ≤ 7 were calculated using jellyfish [49]. From these data, a triple (*key, kmer*_*norm, f req*) was computed for each *k*-mer and each transcript, where *key* is a catagorical variable describing presence or absence of the *k*-mer is a given sequence, *f req* is its relative frequency and *kmer_norm* is the normalized count of the *k*-mer. The entries were converted into features using a MultiLabelBinarizer class from scikit-learn [50]) as binarizer.
2. The Fickett score [35], a measure for the coding potential of sequences, was computed with a custom script inspired by [51].
3. Sequence conservation scores were obtained from the UCSC Table Browser (PhastCons30way) and the scores for individual exons were extracted [52,53]. The PhastCons score was used in two ways; once as mean of all the scores across all the nucleotides and as normalized scores after being divided into 5% bins.
4. RNA secondary structure is often better conserved than the sequence. We therefore utilized structural attributes of the sequences, pairing probabilities of the nucleotides were calculated using RNAplfold [36]. Three different windows of lengths 60, 80, 120 were used for the calculation. Each window was divided into 20 bins, binary encoding each position that falls into the bin with 1 and the rest of the positions with 0.

Since at least a fraction of the annotated lncRNAs is annotated from incomplete transcript models [54], we did not use parameters such as the number of exons, the transcript length, or polyadenylation in any of the classification tasks.

### 4.4. Machine Learning and evaluation framework

We employ a mixture of supervised and unsupervised machine learning techniques, random forests to help us understand the weight of selected features as well as unsupervised techniques which may pick up on features which escape our human perception. Scikit-learn [50,55] was used to perform training, testing and cross-validation of results.

#### Supervised Machine Learning

Random forests are decision tree based classifiers, which work on the basis of an aggregator, training on different subspaces of the training data in parallel. A major advantage of random forest approaches is that they enable us to rank features based an their importance for classification. This allows us to investigate the latter in detail or discard less useful ones to increase training accuracy and time. We used 100 trees for the initial classification, at default settings. 10-fold and 5-fold cross validation was performed with tree sizes [100, 300, 500, 1000], both impurity criteria (gini and entropy), bootstrapping, and taking out of bag samples. A random forest classifier has been trained considering all possible parameter combinations to achieve hyperparameter optimization using grid search. For each combination of feature and data set, we chose the best perfoming hyperparameter setting (details can be found in supl.tab.3). Only little differences in prediction accuracies were observed between training runs on the available parameter space. This is likely related to the rather small size of available datasets, should, however, not diminish gained insights.

#### Unsupervised Machine Learning

As an alternative to the supervised classification strategies discussed above we pursued unsupervised clustering. To this end we used the same four datasets and the same features. To obtain a visual overview of the data we used Principal Component Analysis on the *k*-mer vectors as implemented in R. We then used *k*-means clustering in attempt to distinguish the sequences into three distinct clusters [56]. We further evaluated a convolutional neural network (CNN) based autoencoder instead of a classical clustering approach. For this purpose the *Keras* package (https://keras.io/) was used within the *TensorFlow* framework [57]. We used accuracy, F-measure, and rand index to evaluate the quality of the clusters w.r.t. to distinguishing the three classes of lncRNAs.

### 4.5. Availability

Code used in this publication can be found at github, data is available here.

## Author Contributions

Conceptualization P.F.S., M.E.M.T.W.; supervision P.F.S and J.F., data curation and computational analysis, R.S.; interpretation of results and writing, J.F., R.S., and P.F.S.

## Acknowledgments

We thank Stephanie Kehr for insightful discussions and her advice on all things snoRNA.

## Abbreviations

The following abbreviations are used in this manuscript:

miRNA: microRNA
snoRNA: small nucleolar RNAs
lnRNA: long non-coding RNA
MIRHG: miRNA host gene
SNHG: snoRNA host gene
NoHG: lncRNAs that harbor neither snoRNAs nor miRNAs

