## Supplementary Tables for "Supervised and Unsupervised Classification of lncRNA Subtypes"

**Table 1.** Supplemental Table S1: Results for $k$-means clustering. The *Training Accuracy* column shows the accuracy on the training set, with the count of accurately predicted labels in the *Correctly predicted* column. We show the frequencies of each class in both the training set and the test set in the top row in the following two columns, respectively, with the actual predicted labels in the bottom row. The datasets were split 80-20.

| | Training Accuracy | Correctly predicted (Total) | Labels in trainingset (lnc, mir, sno)<br>Predicted labels [lnc, mir, sno] | Labels in testset (lnc, mir, sno)<br>Predicted labels [lnc, mir, sno] |
|---|---|---|---|---|
| Dataset 1 | 34.42% | 285 (828) | (282, 277, 269)<br>[278,272,278] | (63, 68, 76)<br>[0, 0, 207] |
| Dataset 2 | 33.33% | 579 (1656) | (548, 563, 545)<br>[549,557,550] | (142,127,145)<br>[414, 0, 0] |
| Dataset 3 | 35.76% | 398 (1113) | (371, 373, 369)<br>[371,372,370] | (93, 91, 95)<br>[0, 0, 279] |
| Dataset 4 | 34.79% | 135 (388) | (125, 136, 127)<br>[130, 129, 129] | (37, 26, 35)<br>[98, 0, 0] |

**Table 2.** Supplemental Table S2: Five most important features for classification according to the random forest classifier.. *x-zmers* should be interpreted as all kmers from length x through z.

| Dataset | Featureset | Fickett Score | Rank 1 | Rank 2 | Rank 3 | Rank 4 | Rank 5 |
|---|---|---|---|---|---|---|---|
| Dataset 1 | kmer | without | (3-4mers) | (CG, 3-5mers) | (3-4mers) | (3-6mers) | (3-5mers) |
| | | with | (2-4mers) | (2mers, 4-5mers) | (3-4mers) | (3-4mers) | (4-5mers) |
| | +structure | without | (4-5mers) | (4-5mers) | (4-5mers) | (4-5mers) | (4mers) |
| | | with | (3-4mers) | (4mers) | (4-5mers) | (4-6mers) | (4-6mers) |
| | +conservation | without | (CCG, 4-6mers) | (3-4mers, AGCGG) | (CG, 3-5mers) | (CG, 3-7mers) | |
| | | with | (3-4mers, AGCGG) | (CG, 3-7mers) | (CGA, 4-5mers, 7mers) | (3-4mers, CAGCG) | (CG, 3-7mers) |
| Dataset 2 | kmer | without | GC content | Sequence | (2mers) | (2mers) | (2mers) |
| | | with | Sequence | GC content | Fickett score | (2mers) | (2mers) |
| | +structure | without | GC content | Structure | Structure | Sequence | Structure |
| | | with | GC content | Structure | Fickett score | Structure | Structure |
| | +conservation | without | Conservation | GC content | Sequence | Conservation | Structure |
| | | with | Conservation | GC content | Conservation | Structure | Sequence |
| Dataset 3 | kmer | without | GC content | (7mers) | Sequence | (3-4mers) | (3-4mers) |
| | | with | GC content | (7mers) | Fickett score | Sequence | (7mers) |
| | +structure | without | GC content | (7mers) | Structure | Structure | Structure |
| | | with | GC content | (7mers) | (7mers) | Structure | Structure |
| | +conservation | without | GC content | (7mers) | (3-4mers) | Structure | Structure |
| | | with | (7mers) | GC content | Structure | (7mers) | Structure |
| Dataset 4 | kmer | without | GC content | Sequence | (2mers) | (2mers) | (2-3mers, AAAAA, AAAAAA) |
| | | with | GC content | Fickett score | Sequence | (2mers) | (2-3mers, AAAC, 5-6mers) |
| | +structure | without | GC content | Structure | Structure | Structure | Structure |
| | | with | Structure | GC content | Structure | Structure | Sequence |
| | +conservation | without | GC content | Structure | Sequence | Structure | Structure |
| | | with | GC content | Structure | Structure | Structure | Structure |

**Table 3.** Accuracy after hyperparameter tuning by grid search using 10-fold cross validation metric for supervised machine learning. Indicated are accuracy per feature set and parameters for each feature set as [criterion, number of trees]. Bootstrap and out-of-bag score was always enabled.

| Dataset | Fickett Score | kmer | | +structure | | +conservation | |
|---|---|---|---|---|---|---|---|
| Dataset 1 | without | 85.50% | ['gini', 300] | 85.51% | ['entropy', 100] | 84.06% | ['entropy', 1000] |
| sno: 345, mir: 400, lnc: 400 | with | 84.06% | ['gini', 500] | 80.19% | ['gini', 500] | 82.61% | ['entropy', 100] |
| Dataset 2 | without | 44.69% | ['entropy', 1000] | 47.58% | ['entropy', 1000] | 51.69% | ['entropy', 100] |
| sno: 690, mir: 800, lnc: 750 | with | 40.34% | ['entropy', 300] | 43.96% | ['entropy', 500] | 50.72% | ['entropy', 1000] |
| Dataset 3 | without | 66.67% | ['gini', 1000] | 67.03% | ['entropy', 500] | 70.61% | ['gini', 100] |
| sno: 1287, mir: 464, lnc: 2101 | with | 67.38% | ['entropy', 100] | 65.59% | ['entropy', 500] | 70.25% | ['gini', 300] |
| Dataset 4 | without | 42.86% | ['entropy', 1000] | 50.00% | ['gini', 100] | 62.24% | ['gini', 500] |
| sno: 162, mir: 168, lnc: 750 | with | 41.84% | ['entropy', 1000] | 35.71% | ['entropy', 100] | 50.00% | ['entropy', 100] |
