## Supplementary Figures for "Supervised and Unsupervised Classification of lncRNA Subtypes"

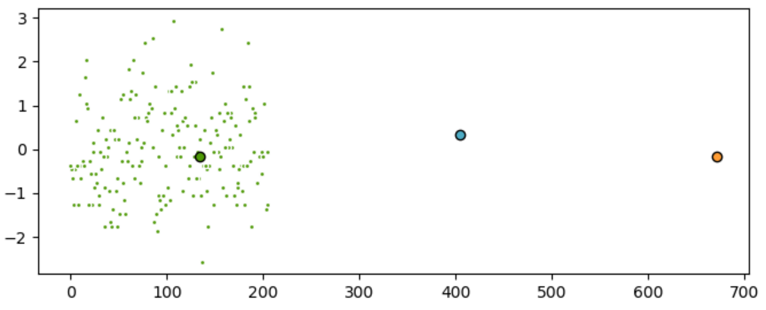


*Figure 1.* $k$-means cluster of the test set of dataset 1. All 207 sequences were clustered as SNOHGs (68 MIRHGs, 76 SNOHGs, 63 NoHGs).

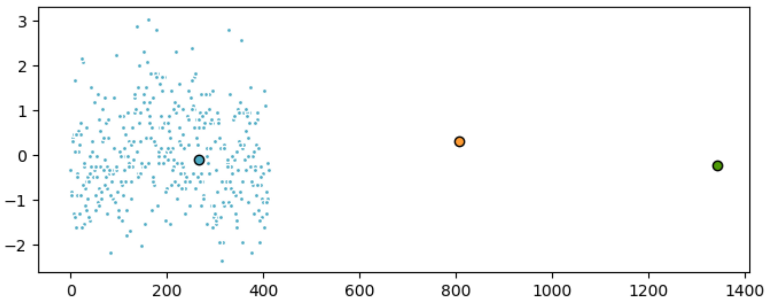


*Figure 2.* $k$-means cluster of the test set of dataset 2. All 414 sequences were clustered as NoHGs (127 MIRHGs, 145 SNOHGs, 142 NoHGs).

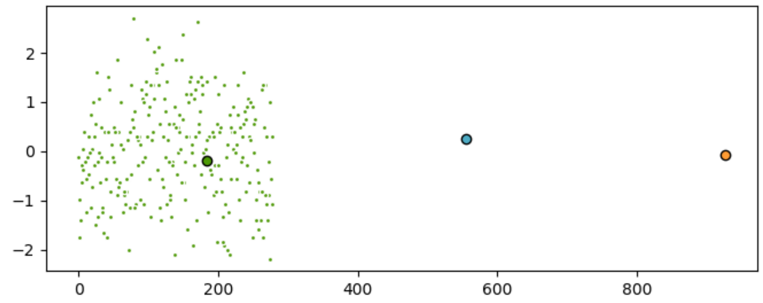


*Figure 3.* $k$-means cluster of the test set of dataset 3. All 279 sequences were clustered as SNOHGs (91 MIRHGs, 95 SNOHGs, 93 NoHGs).

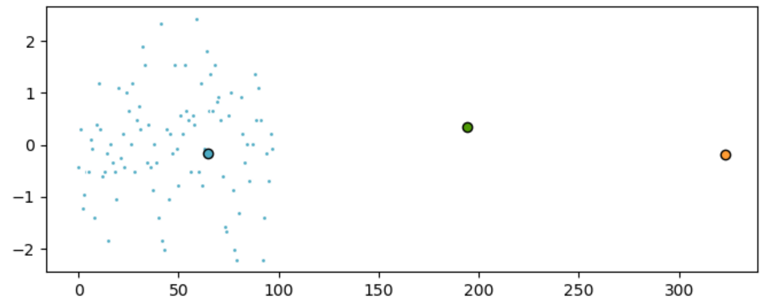


*Figure 4.* $k$-means cluster of the test set of dataset 4. All 98 sequences were clustered as NoHGs (26 MIRHGs, 35 SNOHGs, 37 NoHGs.)

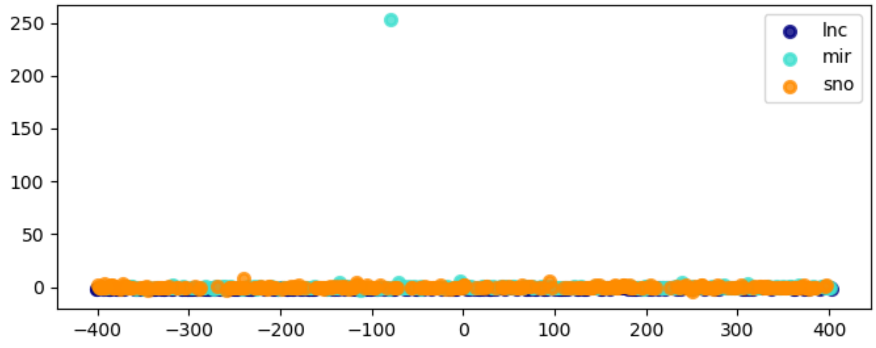


*Figure 5.* PCA of dataset 1. No separation between datasets is possible.

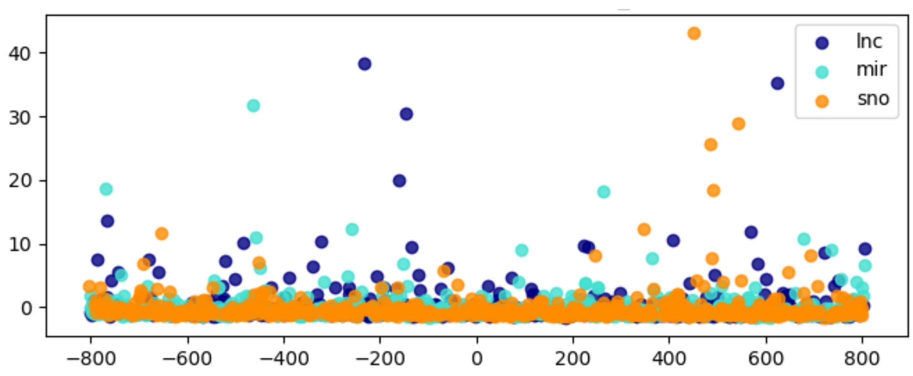


*Figure 6.* PCA of dataset 2. No separation between datasets is possible.

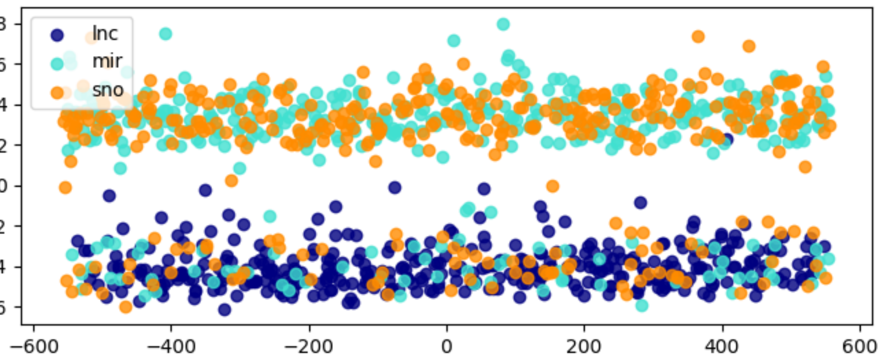


*Figure 7.* PCA of dataset 3. Although NoHGs cluster, no separation between datasets is possible.

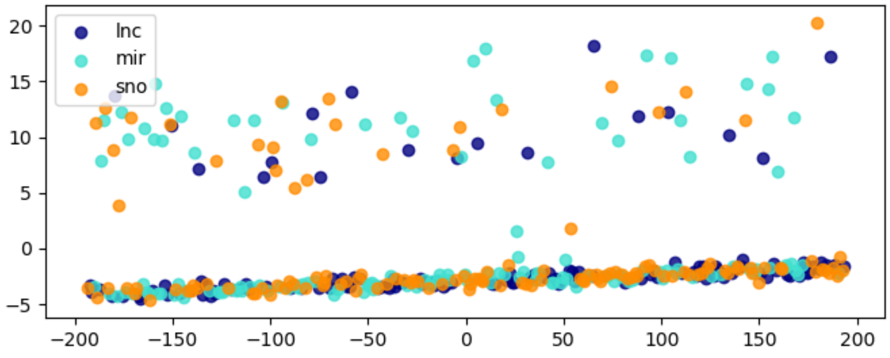


*Figure 8.* PCA of dataset 4. No separation between datasets is possible.
